# Unraveling Viral peptide-G4 Interactions: the NS3 Protease Domain of Yellow Fever Virus Binds G-Quadruplexes with High Specificity and Affinity

**DOI:** 10.64898/2026.03.22.713562

**Authors:** Jiawei Wang, Runfeng Lin, Anne Cucchiarini, Vaclav Brazda, Jean-Louis Mergny

## Abstract

G-quadruplexes (G4s) are critical nucleic acid secondary structures that play pivotal roles in regulating gene expression. In this study, we conducted a proteome-wide *in silico* analysis across multiple viruses causing hemorrhagic fevers to identify candidate proteins containing a conserved G4-binding motif. Four peptides belonging to Marburg, Ebola, Hantaan and Yellow fever viruses were shown to bind to G4 *in vitro*. We selected the NS3 protease domain of Yellow Fever virus for further validation. Biochemical assays demonstrated that the NS3 protease domain binds G4 structures with high specificity and affinity, particularly favoring the parallel conformation. Molecular docking and simulations further revealed that the NS3 protease domain interacts with the terminal G-tetrads and loop regions of G4 via key residues, including PHE40, adopting an insertion and stacking composite binding mode. These findings expand our understanding of virus – G4 interactions and offer novel potential targets for G4-based antiviral strategies.

**Bullet points:**

– We screened viruses causing hemorrhagic fevers for potential G4-binding peptides.
– Four peptides belonging to Marburg, Ebola, Hantaan and Yellow fever viruses were shown to bind to G4 in vitro.
– Biochemical assays demonstrated that the NS3 protease domain of YFV binds G4 structures with high specificity and affinity.

## Introduction

Viruses represent a persistent threat to global health, with their rapid spread and adaptation often resulting in devastating outbreaks. Their ability to exploit diverse transmission routes — ranging from zoonotic spillovers to direct human-to-human contact—poses significant challenges for containment and treatment ^1^. Vector-borne viruses, for instance, leverage ticks, mosquitoes, and other arthropods as efficient transmission vehicles, while others exploit animal reservoirs such as rodents and bats, creating pathways for cross-species transmission ^2-6^. Among these, Viral Hemorrhagic Fevers (VHFs) exemplify the severe impact viruses can have ^7^. Caused by unrelated RNA viruses from families such as Flaviviridae, Bunyaviridae, Filoviridae, and Arenaviridae (see Supplementary Table S1), VHFs – including well-known examples like Ebola and Marburg – often lead to life-threatening illnesses characterized by high mortality, fever, and bleeding disorders ^8^. Transmission via bodily fluids or contact with infected vectors further compounds their public health impact ^9^. These characteristics underscore the need for advanced strategies to understand and combat viral mechanisms.

Despite advances in vaccine development, effective treatments remain elusive for most viruses, in part due to their complex replication cycles and interactions with the host cellular machinery ^10^. G-quadruplexes (G4s), formed by two or more layers of G-tetrads which contain four guanines, are widely distributed in the genetic material of various organisms. These secondary structures have been identified in a wide range of viruses, including those causing VHFs ^11-13^. Flaviviridae viruses (such as Yellow Fever, Dengue, Zika) contain positive-sense RNA genomes in which computational analyses have identified numerous G4-prone sequences. For example, the Zika virus was found to harbor over 60 potential G4 motifs, with several conserved among flaviviruses. Stabilizing these G4 structures with small molecules can suppress viral replication, suggesting G4s in the flavivirus genome act as cis-acting regulatory elements ^14^. Bunyaviridae, which include hantaviruses (*e*.*g*. Hantaan virus) and nairoviruses (*e*.*g*. Crimean-Congo hemorrhagic fever virus), have segmented negative-strand RNAs; while less studied, bioinformatic surveys indicate these genomes harbor putative G4 sequences. It is postulated that during the transcription of these segments or in their complementary RNA, G4 formation could pause the viral polymerase or modulate RNA interactions, thereby influencing gene expression ^15^. Filoviridae (*e*.*g*. Ebola and Marburg viruses) likewise feature G-rich tracts in their negative-sense RNA genomes. The first evidence of an RNA G4 in a negative-strand virus was demonstrated in the L gene of Ebola virus: a conserved G4 RNA motif whose stabilization by a G4 ligand markedly reduced expression of the viral polymerase and inhibited viral genome replication in a mini-replicon system ^16^. Similarly, Arenaviridae (*e*.*g*., Lassa virus, with bi-segmented negative RNA) contain G-rich regions – for instance, the intergenic regions or non-coding tails – that may fold into G4. Although direct experimental validation in Bunyaviridae and Arenaviridae is still lacking, the conservation of G4 motifs in these viral genomes across strains hints at functional importance ^15^.

A growing area of interest is the interaction between viral proteins and host nucleic acid structures, particularly G4, as their exploitation by viruses may play a pivotal role in viral replication and gene expression ^17^. There is no shortage of proteins with which to interact during all of these processes. Investigating whether viruses encode G4-binding proteins (G4BPs) offers a promising avenue to uncover the molecular underpinnings of viral pathogenesis. Such interaction sites may constitute novel therapeutic targets, as recently illustrated by the SUD domain of the Nsp3 SARS-CoV-2 protein ^18-19^. The arginine-glycine or arginine-glycine-glycine (RG/RGG) motif is a typical class of domain among G4BPs ^20^. Originally referred to as the RGG box, this motif enables a protein to bind to double-stranded mRNA molecules ^21^. It was then discovered that Fragile X mental retardation protein (FMRP) binding to G-rich RNAs *in vitro* relies solely on the RGG motif, which specifically interacts with both natural and in vitro-selected G4-containing RNAs ^22^. Subsequently, numerous proteins with RGG motif were identified to interact with G4 structures, including Nucleolin via its RGG domain ^23^ or hnRNPA1 ^24^.

Although the overall RGG domains have been extensively studied, the specific role of the RGG primary sequence remains unclear. To address this, Huang *et al*. ^25^ systematically examined the binding affinity and interaction mechanisms between RGG-motif peptides and G4 structures. Arginines and phenylalanines at precise positions within the RGG motif may enhance binding to G4 by facilitating additional hydrogen bonding and π-stacking interactions with nucleobases. In the process, a new G4 DNA-binding protein was also identified, the cold-inducible RNA-binding protein (CIRBP). Simultaneously, Brázda *et al*. ^26^ attempted to understand the binding mode of RGG motifs to G4 from another perspective by means of statistical generalization. 77 described G4BPs of *Homo sapiens* were dealt with Cluster analysis with bootstrap resampling, revealing similarities and differences in amino acid composition of specific G4BPs. A 20-amino-acid long motif (NIQI, for *Novel Interesting Quadruplex Interaction*), RGRGRGRGGGSGGSGGRGRG, was defined and found to be common to all characterized G4BPs analyzed. Based on the NIQI motif, Červe ň *et al*. ^27^ predicted G4BPs in plants with the help of Find Individual Motif Occurrences (FIMO) approach and validated these predictions with molecular docking.

However, whether viruses causing hemorrhagic fevers encode their own G4-binding proteins remains unclear, representing a critical gap in our understanding of virus – host interactions. In this work, we address this question by combining bioinformatic motif scanning with experimental validation to identify and characterize viral G4-binding proteins from VHFs. Using the conserved G4-recognition motif NIQI as a search probe, we screened the proteomes of several pathogenic viruses and uncovered multiple candidate G4-binding sequences. 7 candidate peptides were then synthesized and tested for G4 recognition, providing experimental confirmation of this interaction for four of these peptides. We then demonstrated that one of these candidates – located in the Yellow Fever virus NS3 protein – indeed exhibits robust and specific binding to G4. Through biochemical assays and structural modeling, we further elucidate the mode of interaction between this viral protein and G4 structures, revealing key residues that engage the G4 motif. Together, our findings fill a notable knowledge gap by providing the evidence that a non-structural protein of Yellow Fever Virus (YFV) can directly recognize a G4.

This insight not only deepens our understanding of how viruses may exploit host nucleic acid structures, but also highlights G4 elements as potential new targets for antiviral intervention.

### Experimental Section

#### Bioinformatics

Identification of G4-binding peptides in virus was performed using FIMO (https://meme-suite.org/meme/tools/fimo) ^28^, with a threshold of *p* = 0.001 to minimize false-positive results. As the input for motif scanning, we separately used 7 reference proteome sequences for CCHFV, Ebola, Hantaan, Lassa, Marburg, Nipah and Yellow fever. Motif scanning was carried out for the NIQI motif.

#### Samples

##### Oligonucleotides

All the oligonucleotides were purchased from Eurogentec (Belgium) in the form of dried powder and dissolved in distilled and deionized water. DNA and RNA sequences were purified through RP cartridge and HPLC-RP methods, respectively. Primary sequences are shown in Table 1. All the samples were stored at -20℃.

**Table 1.**
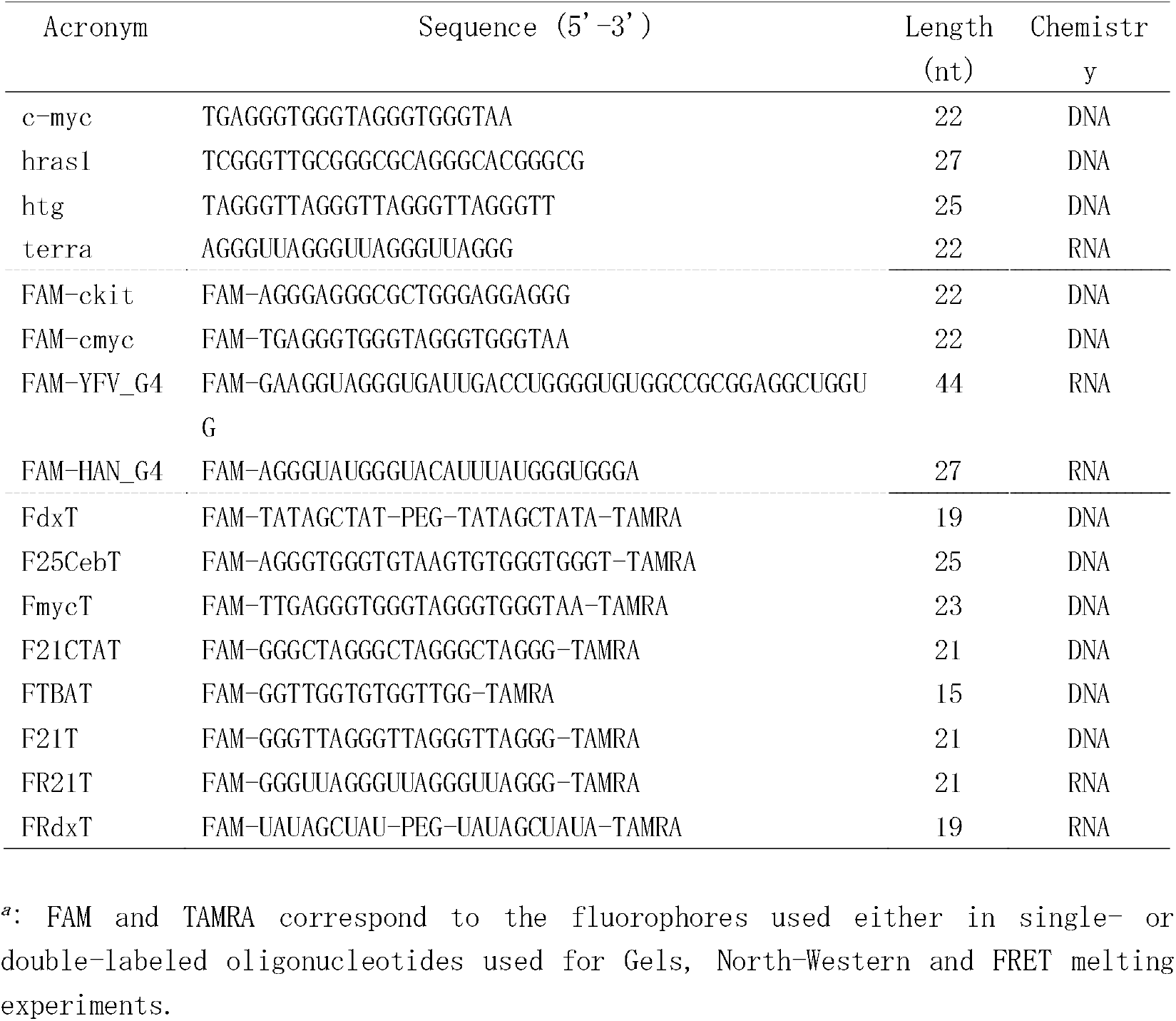
Oligonucleotide sequences^*a*^.

##### Peptides

All peptides were purchased from Genecust (France) in the form of dried powder and dissolved in distilled and deionized water. The peptides are acetylated at N-terminal and amidated at C-terminal and delivered with a final purification of >95%. Peptide sequences are shown in Table 2. All samples were stored at -20℃.

**Table 2.**
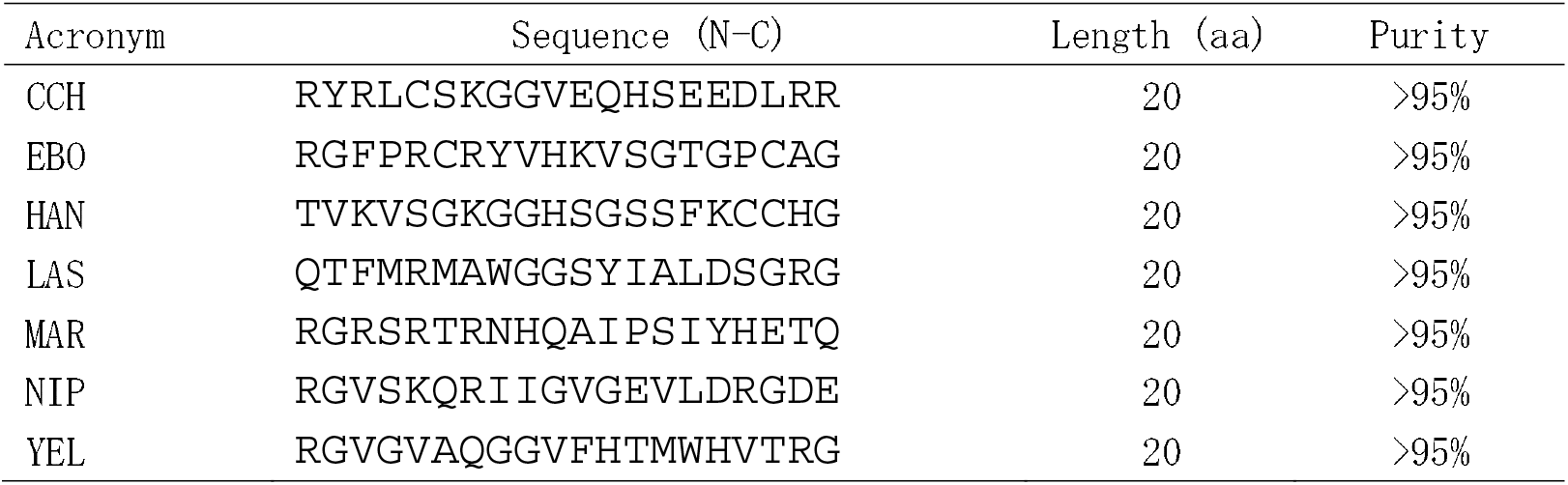
Viral peptides tested here.

##### Plasmids

All plasmids were purchased from Twist Bioscience (USA) in the form of dried powder and dissolved in distilled and deionized water. More details regarding these plasmids are shown in Table S2. All samples were stored at -20℃.

#### Non denaturing Electrophoresis

20 μl solution included 0.25 μM FAM-ckit, 20 mM HEPES (pH = 7.2) and 100 mM KCl with 0, 0.05, 0.125, 0.25, 0.5, 1.25 or 2.5 μM peptide. The samples without peptides were first annealed at 95℃ for 5 min. After slowly cooling down to room temperature, peptides were added at different concentrations. The samples with 4 *μ*l 60% sucrose were loaded on a 15% polyacrylamide gel (acrylamide: bisacrylamide = 37.5: 1) prepared in a 1X Tris-borate-EDTA (TBE) buffer. Electrophoresis was carried out for 210 min at 60V, at 4℃, in 1X TBE buffer containing 20 mM KCl.

#### FRET-melting

25 *μ*l solution included 0.2 *μ*M double-fluorophore-labeled pre-annealed sequences, 10 mM LiCaco buffer (pH = 7.2) containing 10 mM KCl and 90 mM LiCl with 0, 0.2, 1 or 5 *μ* M peptide. Samples were placed in 96-well plates and FRET-melting was performed in a qPCR machine to monitor the change of the donor fluorescence over time, as described previously ^29-31^. The temperature gradient was 1℃/min, between 20℃ and 95℃.

#### Circular dichroism (CD) & Thermal differential spectrum (TDS)

CD spectra were recorded with a J-1500 spectropolarimeter (Jasco) from 350 nm to 220 nm at 25℃. The scan rate was 200 nm/min and the spectra were averaged by two accumulations. TDS correspond to the difference between the UV-absorbance spectra (between 350 nm and 220 nm) at 95℃ and 5℃ on a Jasco spectrophotomer through the UV channel ^32^.

#### Protein expression and purification

The plasmids encoding the two protein constructs were cloned by heat-shock transformation method in *E. coli* first. After extracting the corresponding plasmids, His-fused YFV_pro and His-fused HAN_pro constructs were both expressed in Rosetta (DE3) strain cells. The pET-28a-transformed cells were cultured in LB buffer with 30 μg/ml Kanamycin and 20 μg/ml Chloramphenicol, grown at 37℃ until OD_600 nm_ reached 0.7, then induced with 0.5 mM IPTG. Different temperatures (16, 25, and 37° C) were tested for growth for some time (overnight, overnight, and 3h, respectively). The cells were harvested by centrifugation at 6000 rpm for 30 min at 4℃. After resuspending the pellets with deionized water in 50 ml Falcon tubes, the cells were centrifuged at 5000 rpm for 20 min at 4℃. The pellets were stored at -20℃ before purification.

10 mL of 30 mM HEPES buffer containing 150 mM NaCl, one protease inhibitor cocktail tablet, and 1 mg/mL lysozyme was added to the pellet. The mixture was resuspended by vortexing and incubated on ice for 20 min. The suspension was subjected to sonication on ice for 8 cycles, alternating 20 s of pulse and 60 s of resting. The lysate was then centrifuged at 13,000 rpm for 30 min at 4°C, and the supernatant was discarded. The pellet was resuspended in 30 mM HEPES buffer containing 8 M urea and 150 mM NaCl, followed by a second round of sonication on ice for 5 cycles, alternating 20 s of pulse and 60 s of resting. Finally, the sample was centrifuged at 13,000 rpm for 10 minutes at 4°C, and the supernatant was collected.

GE AKTA pure was used for the purification. The sample was loaded on a pre-equilibrated 5 mL Ni-column, which was re-equilibrated with 10 column volumes (CV) of buffer. Protein renaturation was performed on the column using 15 CV of buffer containing a gradient of 8 M to 0.1 M urea. Subsequently, the target protein was eluted with 10 CV of buffer containing 0.1 M urea, 150 mM NaCl, 30 mM HEPES and a gradient of 0 to 500 mM imidazole. The sample HAN_pro was desalted using a 3 kDa molecular weight cutoff ultrafiltration tube by centrifugation at 5,000 rpm for 30 min at 4°C. The buffer was refilled with imidazole-free buffer and repeated rounds of centrifugation were then performed. In contrast, the YFV_pro was eluted during the renaturation process, meaning that the sample still contained a high concentration of urea. To address this, urea was gradually removed through dialysis, with the final concentration maintained at 2M without causing precipitation. Finally, glycerol was added to the samples to a final concentration of 10%, and the samples were stored at -20°C. According to our calculations, the final urea concentration in follow-up experiments should remain within an acceptance range (<0.5 M) and if needed, control groups containing urea would be set up to ensure that it would have little effect.

#### Western blotting

The sample was mixed with 2×Laemmli sample buffer at a 1:1 ratio, then immediately denaturing at 95℃ for 10 min. Loading the sample on a 12% SDS-polyacrylamide gel and running the gel at 120 V, 4℃ for 90 min. The proteins were transferred on a NC membrane which was then blocked with TBST buffer containing 5% skim milk at room temperature for 1 h. Subsequently, the membrane was incubated with a primary antibody (Anti-His mouse) at room temperature for 1 h. After rinsing with TBST buffer the membrane was incubated with the secondary antibody (IRDye 800CW goat anti-mouse) at 4℃ overnight, and finally imaged with a ChemiDoc system through the IR800 channel.

#### South-Western blotting and North-Western blotting

The sample was mixed with 2×Laemmli sample buffer in a 1:1 ratio, then immediately denaturing at 95 ℃ for 10 min. Next, the sample was loaded on a 12% SDS-polyacrylamide gel and the gel was run at 120 V, 4℃ for 90 min. The proteins were transferred to a Nitrocellulose membrane which was then renatured at 4℃ for 2 h with the buffer containing 20 mM Tris-HCl (pH 7.5), 1mM EDTA, 10% glycerol and 0.1% Triton X-100. Next, the membrane was blocked with TBST buffer containing 5% skim milk at room temperature for 1 h. Subsequently, the membrane was hybridized with FAM-labeled DNA G4 (dG4) or RNA G4 (rG4) at 4℃ overnight, and finally imaged with a ChemiDoc system through the Alexa488 channel.

#### Thioflavin-T competition assay

Thioflavin-T (ThT) may be used as a fluorescent light-up probe for G4 formation ^33^. Mixing 0.1 *μ*M pre-annealed G4s of different topologies with ThT in a ratio of 1:1. Then 0, 5, 10, 20, 50, 100, 200, 500, 1000 or 2000 nM proteins were added to the samples which were placed into a 96-well plate. The fluorescence intensity was measured with a qPCR machine at room temperature. Each sample was tested in duplicate and the results were averaged.

#### Fluorescence anisotropy

0.1 *μ*M pre-annealed FAM-labeled G4 was mixed with 0, 10, 20, 50, 100, 200, 500, 1000 or 2000 nM proteins. A Varian Cary Eclipse fluorescence spectrometer (Ex=492 nm, Em=510-600 nm) was used to measure the intensities at vertical (I_VV_) and horizontal (I_VH_) polarizations.

The anisotropy (r) is calculated using the formula:

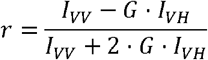

The G-factor accounts for detector sensitivity and is calculated as:

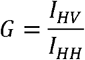

Fitting r as a function of protein concentration and the binding affinity of protein with G4 was calculated as K_d_.

#### Structural simulation and Docking

AlphaFold3 (https://alphafoldserver.com/) was used to simulate the structure of truncated protein YFV_pro and we then performed molecular docking of this protein with a parallel G4 (c-myc; PDB: 1XAV) by HADDOCK (https://rascar.science.uu.nl/haddock2.4/).

#### Molecular dynamics

MD simulations were performed using AMBER24 ^34^ with the OL21 ^35^ force field for nucleic acids and ff19SB ^36^ for proteins to generate the G4–protein complex topology. The system was neutralized with 19 Na^+^/K^+^ counterions and solvated using the TIP3P water model ^37^. Energy minimization was performed using a two-stage protocol: a total of 50,000 minimization cycles were executed (with the first 25,000 cycles employing the steepest descent algorithm, followed by conjugate gradient minimization). The system was then heated under constant volume (NVT) conditions over 50,000 steps (with a time step of 0.002 ps) using a stepped temperature ramp from 0 to 300 K (first 20,000 steps) and maintained at 300 K thereafter, with Langevin dynamics and a collision frequency of 2 (ntt=3, gamma_ln=2.0). Next, volume equilibration was carried out under constant pressure (NPT) conditions at 1 atm and 300 K for 25,000,000 steps (dt=0.002 ps) and a nonbonded cutoff of 14.0 Å, with Particle Mesh Ewald (PME) settings (FFT grid dimensions of 64 and order 4) ensuring accurate treatment of long-range electrostatics. Finally, production simulations were run in the NVT ensemble for 525,000,000 steps (equivalent to 1.05 µs, dt=0.002 ps) at 300 K using the Langevin thermostat with collision frequency of 1 (ntt=3, gamma_ln=1.0) and the same PME parameters, with trajectory data recorded every 50,000 steps. The agglomerative clustering algorithm with complete linkage and a distance threshold of 3 Å enforced maximum structural deviations below 3 Å within the same clusters ^38^. The MD trajectories were analyzed using MDTraj ^39^, and the interface residues were defined based on the buried surface area (BSA) calculated by the solvent-accessible surface area (SASA) difference between the complex and the isolated chain ^40^:

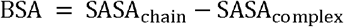

The similarity of the interface is defined by the Jaccard index between the predicted structure interface residues and the stabilized structure interface residues (after 850 ns), which is:

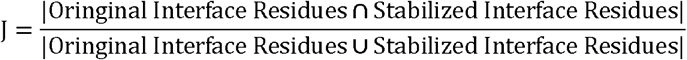

Molecular structures were displayed using diverse visualization styles, with color gradients applied to depict temporal progression. Trajectory playback and RMSD overlays were presented, while clustering algorithms were utilized to identify distinct conformational states. The visualization allowed for interactive exploration, including zooming, rotating, and translating the molecular structures. Key frames were exported as PDB files for further analysis.

## Results

### Identification of G4-binding peptides using the FIMO approach

The matching results showed that each viral proteome contains several sequences with different extents of similarity. All predictions with *p*-values less than 0.001 were sorted by the score and categorized by virus species in Supplementary Table S3. We selected the best rated candidate (with the lowest *p*-value) for each virus and finally listed 7 peptide candidates (CCH, EBO, HAN, LAS, MAR, NIP, YEL) (Table 2).

### Experimental confirmation of G4 recognition by viral peptides

After obtaining the 7 candidate peptides, we first performed an electrophoresis assay to test G4-peptide interactions. If a peptide binds to the pre-annealed FAM-labeled fluorescent G4, one would either expect a shift in migration in a non-denaturing gel (the complex would migrate slower than the band of G4 alone) or the disappearance of the G4 band (if the interaction quenches the fluorophore or leads to aggregates). Upon increasing peptide concentration, when the ratio of FAM-G4 to peptide reached 1:2, some bands (EBO, HAN, MAR and YEL) began to weaken (Fig. 1a). For a ratio of 1:10, some bands almost completely disappeared.

**Figure 1.**
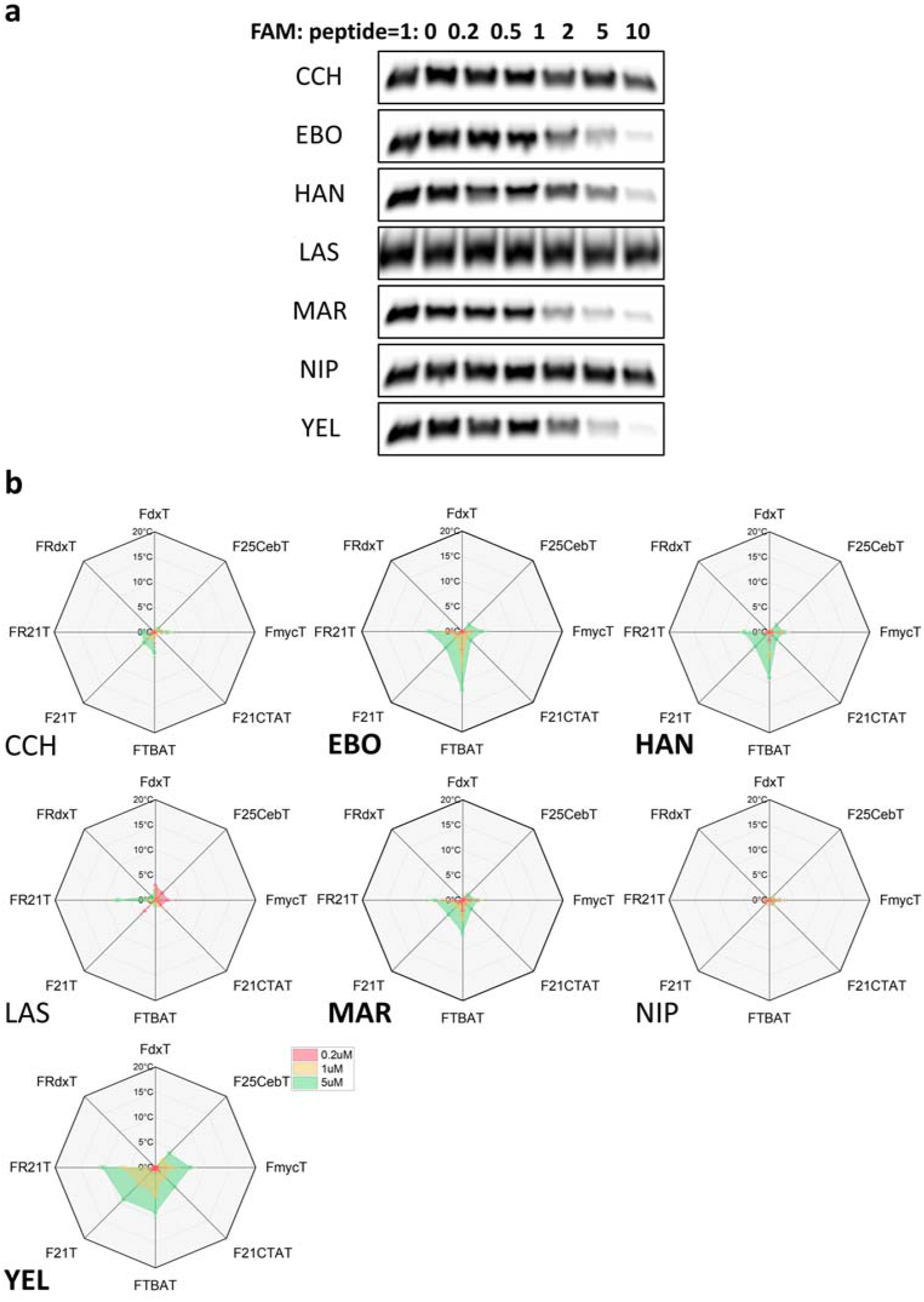
**(a)** Electrophoretic graphs of FAM-ckit interacting with virus peptides at different concentrations (RHAU peptide shown in Fig. S1). **(b)** Radar charts of differences in T_m_ upon binding of virus peptides at three different concentrations to 8 different nucleic acid secondary structures followed by FRET melting. FdxT: hairpin duplex; F25CebT, FmycT: parallel G4s; F21CTAT, FTBAT: antiparallel G4s; F21T: hybrid G4; FR21T: parallel rG4; FRdxT: RNA hairpin duplex.

Next, we performed FRET-melting experiments ^41^ to confirm the binding of peptides to G4. The oligonucleotides used here were double-labeled, with a FAM modification at the 5’-end and TAMRA group at the 3’-end, allowing resonance energy transfer from FAM to TAMRA thanks to their spatial proximity induced by the formation of a secondary structure (hairpin or quadruplex). As the temperature rises, the fluorescence of FAM is restored when the sequence unfolds; thus, we can determine the melting temperature (T_m_) of the corresponding structure. Binding of the peptide to the G4 should stabilize the pre-folded structure and increase its T_m_. The change in T_m_ (ΔT_m_) upon addition of the peptide provides a rough estimate of the binding strength of the peptide to different secondary structures. We chose a set of nucleic acid structures involving DNA and RNA hairpin duplexes (FdxT & FRdxT, respectively) and a variety of dG4 and rG4 conformations, involving parallel, antiparallel, or hybrid quadruplexes. Results are summarized in Fig. 1b and representative melting curves are provided in Fig. S2. No significant stabilization of the duplex was found for any of the peptides, except LAS, while four peptides (EBO, HAN, MAR, YEL) stabilize the different G4s to various extents, demonstrating that these peptides exhibited preferential binding to G4s compared to duplex controls. The presence of these single-strands did not affect ΔT_m_, demonstrating that ssDNA could not act as “decoy” to trap these peptides and prevent stabilization of the G4, demonstrating selective interaction.

Of note, the FRET-melting results (Fig. 1b) are fully consistent with electrophoresis results (Fig. 1a) as the same four candidates were identified by both methods. Among these candidates, YEL exhibited the most pronounced stabilization in terms of ΔT_m_. Among the three other candidates, we selected HAN as the most promising peptide, based on the relatively small size of the minimal main protein domain to which the peptide belongs.

### Expression and purification of the candidate proteins

The observation that individual 20-aa peptides bind to G4s does not imply that the correspond full-length protein or protein domain would also interact with G4s. To check this hypothesis, we identified the complete minimal protein domains in which these two peptides are located (Fig. 2a). The protein to which YEL belongs is Peptidase S7, Flavivirus NS3 serine protease (NCBI: NP_041726). This 152-aa protease is also part of Yellow Fever Non-structural protein NS3 that corresponds to the N-terminal end ^42^, while the C-terminal domain of NS3 functions as an RNA helicase and NTPase, playing a crucial role in genome replication and viral RNA synthesis ^43^.

**Figure 2.**
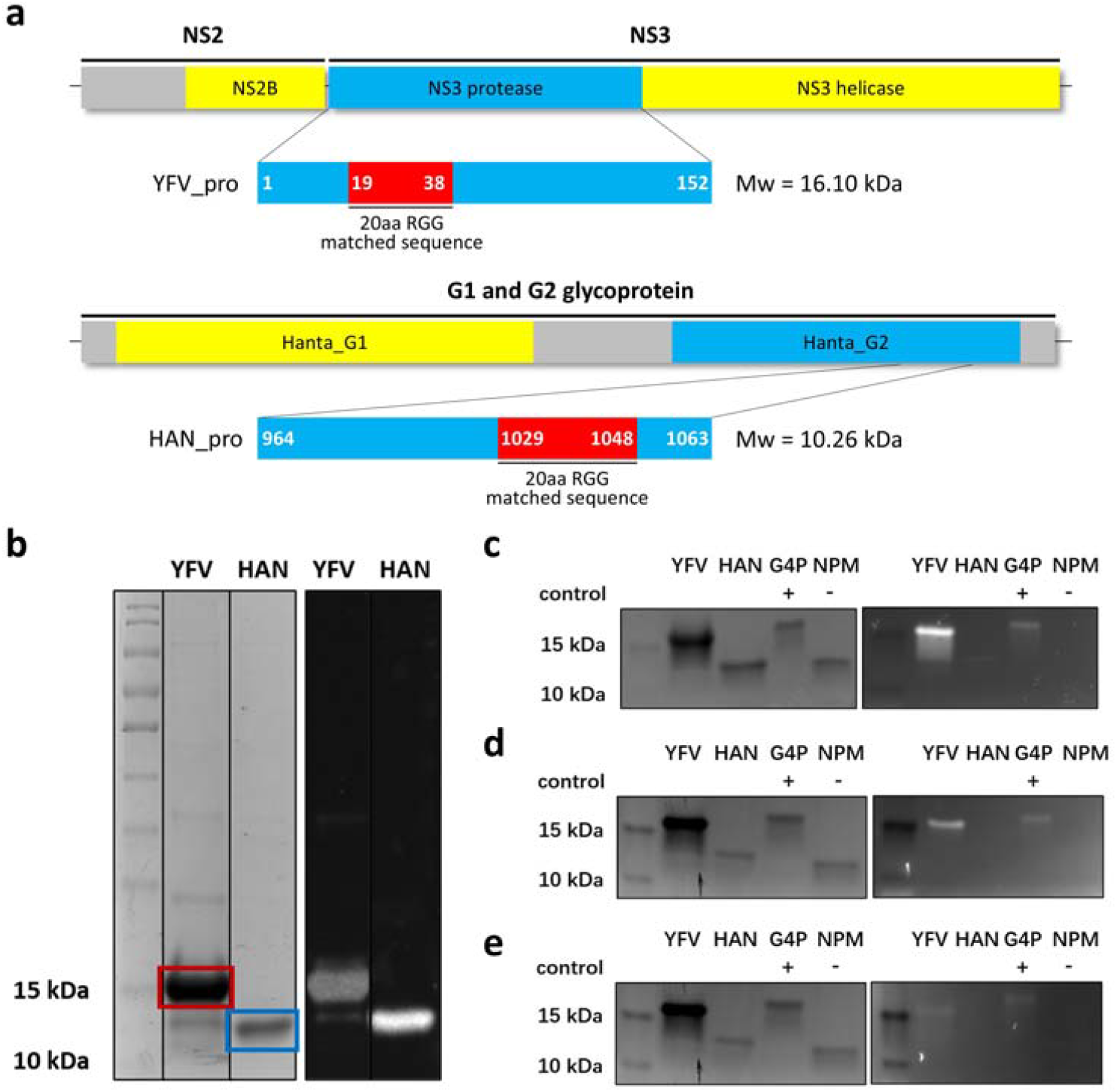
(**a**) Location of the YFV_pro (top) and HAN_pro (down) peptides tested here. (**b**) Coomassie Brilliant Blue staining (left) and Western Blotting (right) of protein YFV_pro and HAN_pro after purification. (**c**) Coomassie Brilliant Blue staining (left) and South-Western Blotting (right) of protein YFV_pro and HAN_pro with DNA probe FAM-ckit. (**d**) Coomassie Brilliant Blue staining (left) and Nouth-Western Blotting (right) of protein YFV_pro and HAN_pro with RNA probe FAM-YFV_G4. (**e**) Coomassie Brilliant Blue staining (left) and Nouth-Western Blotting (right) of protein YFV_pro and HAN_pro with RNA probe FAM-HAN_G4. G4P and NPM are positive and negative peptide controls, (the former is able to recognize G4 structures).

The HAN peptide is located within the G1 and G2 glycoproteins (NCBI: AFM85164) of Orthohantavirus hantanense. Although the G2 glycoprotein alone is 455-aa long, it exhibits clearly distinct domains. We truncated the independent structural domain to 100 aa at the C-terminal end where the HAN peptide is positioned (Fig. S3). To efficiently express the two target proteins, we chose the vector pET-28a and added a His-tag at the N-terminal for subsequent purification. We then cultivated them at 37℃, the optimal temperature for YFV_pro and HAN_pro expression (Fig. S4). After disruption and centrifugation, the supernatant and precipitate were loaded on a gel.

Our result indicated that the target proteins were expressed in inclusion bodies (Fig. S5). At last, samples containing YFV_pro and HAN_pro were obtained by AKTA purification and post-processed. Coomassie Brilliant Blue staining and Western Blotting showed that purification products were pure (Fig. 2b).

### Binding ability of YFV_pro and HAN_pro to G4

Having obtained the purified polypeptides, we next tested their binding to G4s through different methods. First, we performed a South-Western Blotting assay with YFV_pro and HAN_pro using a FAM-labeled dG4. We observed a bright band around 15 kDa for YFV_pro and a weaker one above 10 kDa for HAN_pro (Fig. 2c). We next tried a different fluorescent sequence, choosing rG4 to conduct a North-Western Blotting experiment.

We chose to assay rG4s corresponding to natural sequences found in the genomes of Yellow Fever and Hantaan viruses. The Candidate sequence YFV_G4 was found by Schult *et al*. ^44^, and HAN_G4 was identified with G4-Hunter ^45-46^. Short oligoribonucleotides were chemically synthesized and we next experimentally confirmed rG4 formation through a combination of techniques. First, circular dichroism revealed a negative peak around 240 nm and a positive peak around 265 nm for both sequences, in agreement with rG4 formation (Fig. S6). Thermal Difference Spectra (TDS) confirmed this conclusion, as evidenced by the negative peak at 295 nm and overall shape of these spectra (Fig. S7).

The blotting results are shown in Fig. 2d-e: a clear fluorescent band is observed for the protein YFV_pro, confirming its ability to bind to dG4 and rG4 (G4P, a peptide known to bind to G4s is shown for comparison). Much weaker (Fig 2c, with a fluorescent dG4 probe) or nearly absent signals (Fig 2d-e with fluorescent rG4 probes) were found with HAN_pro, suggesting a lower affinity of HAN_pro for quadruplexes.

We then performed FRET-melting experiments, similar to those obtained with the short peptides (Fig. 3a). The radar charts demonstrate that the intact YFV_pro structural domain stabilize all quadruplexes in a concentration-dependent manner, but has negligible effect on the duplex, confirming its specificity. Interestingly, ΔT_m_ with the full domain were higher than with the short 20aa long YEL peptide (Fig. 1b). For G4 structures, the strongest stabilization at all concentrations was found with the parallel G4 c-myc (up to +25℃),. In contrast, the HAN_pro domain was nearly inactive, except at the highest concentration on FmycT, with a ΔT_m_ of ≈5°C. These observations are in agreement with the low or absent signal in South-Western and North-Western plots, and confirm that HAN_pro is a much weaker G4-binding protein than YFV_pro. Its performance is actually lower than the ones of the HAN 20-aa peptide.

**Figure 3.**
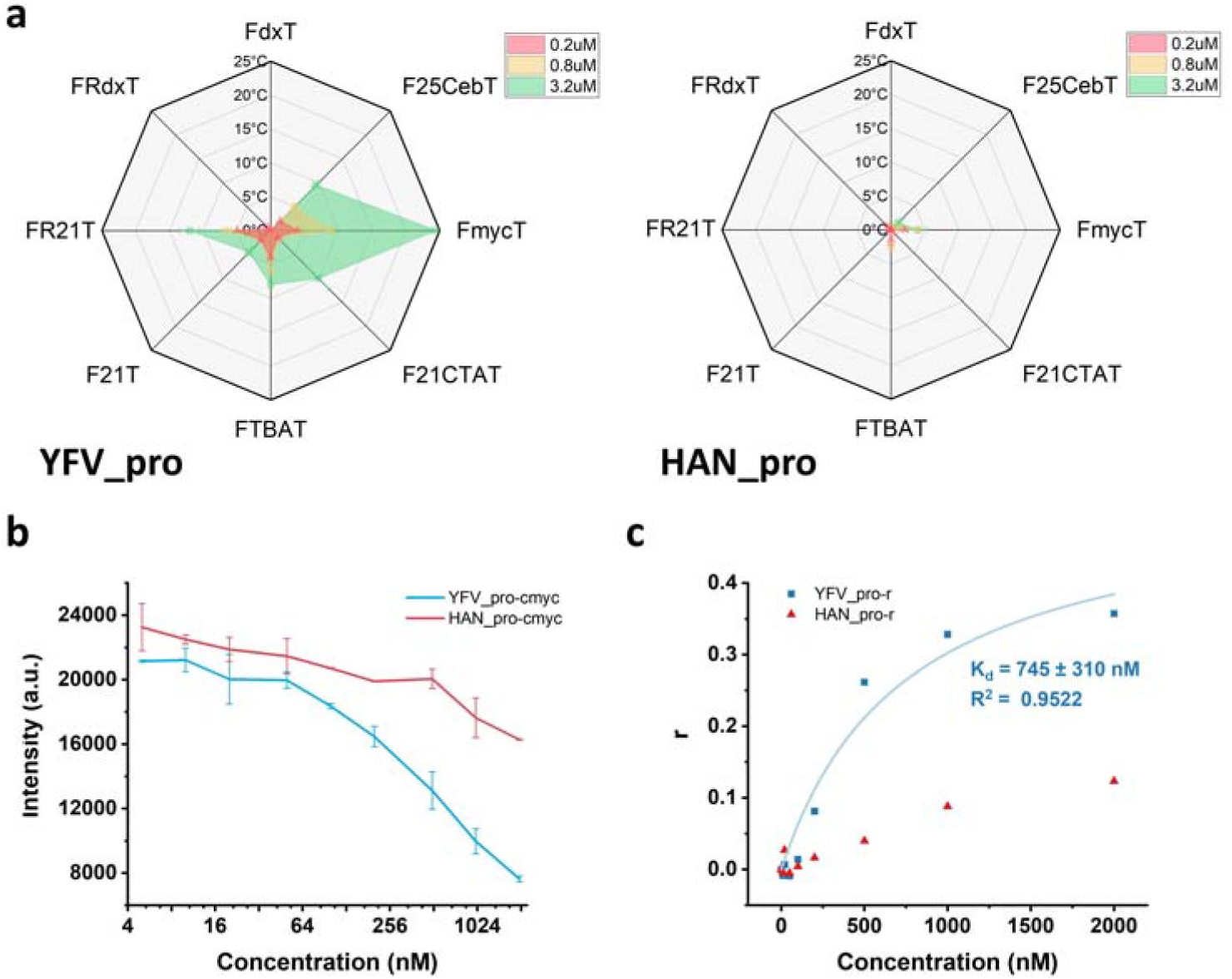
(**a**) Radar charts of ΔT_m_ upon addition of proteins YFV_pro and HAN_pro at 3 different concentrations. Eight different nucleic acid secondary structures were considered: FdxT: duplex; F25CebT, FmycT: parallel G4; F21CTAT, FTBAT: antiparallel G4; F21T: hybrid G4; FR21T: parallel rG4; FRdxT: RNA duplex. (**b**) Fluorescence intensities of ThT with parallel G4 c-myc after addition of YFV_pro and HAN_pro in gradient concentrations. (**c**) Fluorescence anisotropy titration with YFV_pro and HAN_pro, allowing to estimate a dissociation constant.

A third series of experiments was performed to assess binding, based on Thioflavin T (ThT) fluorescent competition assay (Fig. 3b). As a common G4 ligand, ThT emits fluorescence at 490 nm upon excitation with 425 nm after binding to G4 ^33^, and competition should occur upon addition of a G4 ligand. For the YFV_pro domain, the fluorescence intensity of ThT bound to c-myc decreased with increasing protein concentration, confirming competition, with an apparent IC_50_ of 515 nM. For HAN_pro, the competition was less efficient, with a minimal decline in fluorescence. Again, these results illustrate the higher affinity of the YFV_pro for G4, in agreement with the North/South Western and FRET-melting assays.

Last, given that both YFV_pro and HAN_pro bind more strongly to parallel G4 c-myc, we performed Fluorescence Anisotropy titrations (Fig. 3c). When a fluorophore attached to a relatively small molecule (a G4) binds to a larger protein, its tendency to rotate freely tends to decrease, leading to increase in fluorescence anisotropy. Based on changes in r value as a function of YFV_pro concentration, we could calculate a dissociation constant K_d_ of 745 ± 310 nM. No K_d_ could be determined for HAN_pro.

### Molecular docking of YFV_pro with c-myc

To better explore the binding mode between YFV_pro and c-myc, we got a first prediction through molecular docking. Initially, we input the protein YFV_pro and the nucleic acid c-myc, designating the 20-aa RGG matched motif in the protein as interaction sites. The server then generated 13 clusters comprising a total of 52 potential models which were scored and ranked according to its scoring system. During the docking process, a knowledge-based scoring function was used to evaluate the binding energy, and the conformation with the lowest energy was selected as the best prediction. According to the most reliable cluster from HADDOCK, the complex exhibited favorable binding stability, as reflected by the HADDOCK score of -100.2, with significant contributions from van der Waals and electrostatic interactions. The Z-score of -2.2 further supported the reliability of this binding mode, suggesting that this cluster was among the most favorable conformations sampled. The buried surface area (BSA) of 1711.9 Å^2^ suggested a well-packed binding interface, which was indicative of strong intermolecular interactions. Furthermore, the low RMSD value of 0.9 Å highlighted the structural consistency within the cluster, reinforcing the robustness of the predicted binding pose. However, the relatively high restraint violation energy of 236.6 suggested some deviations from the experimental constraints, which may indicate potential structural discrepancies that required further investigation. Additionally, the desolvation energy contribution was relatively low, implying that portions of the binding interface may remain exposed to the solvent, potentially affecting the overall stability of the complex.

At the same time, from the model we could see that GLN 10 and ARG 37 interact with the 5-terminal G-quartet plane, such as H-bonds (Fig. S8). Several residues, including THR 12, PHE 13, PHE 40, and LYS 47 are found in close proximity to the lateral loops of the G4 structure, suggesting a loop-insertion mode. dT1, located in the flanking regions of the G4, appears to be engaged in direct protein interactions. These nucleotides may be stabilized within a protein pocket, suggesting that the protein can recognize not only the core G4 structure but also its surrounding sequence context. It showed that there was a strong binding between these two elements, while we still expected a dynamic stability of the complex.

### MD simulation of docking result

To further validate these findings, we performed molecular dynamics simulations to assess the dynamic stability of the complex, while additional analyses of key interacting residues may provide deeper insights into the binding mechanism. MD simulations were performed for 1.05□**μ**s to examine the predicted stability of the complex by measuring the overall structure RMSD and the similarity of the interface. The global backbone RMSD fluctuates relative to the original structure and has reached an average above 3.5□**Å**, which suggests that the complex has adjusted into other conformations (Fig. S9). In addition, the interface is more flexible than the whole complex, likely due to DNA-induced flexibility. For example, residues near the DNA groove often undergo dynamic rearrangements to accommodate sequence-specific interactions, as observed in studies of transcription factors like PurR ^47-48^. To assess whether the complex had stabilized, the trajectories were clustered based on the pairwise RMSD every ns. It is obvious that this complex is very flexible and only becomes stabilized after approximately 850□ns, at which point none of the structural pairs have an RMSD above 3□**Å**. By measuring the RMSD and RMSF (Fig. S10) relative to the 850□ns structure for frames after this point, we observe a very stable complex with a mean RMSD of 1.59 ± 0.16□**Å**. Additionally, the RMSD of the protein and DNA were 1.40 ± 0.12□**Å** and 1.47 ± 0.21□**Å**, respectively, suggesting that both the protein and DNA structures are quite rigid ^49^. The Jaccard index (Fig. S11) of the interface between the original predicted structure and the stabilized structure is around 0.7, which indicates a highly similar and stable interface, with the main difference arising from the insertion of the DNA 5’ end into the protein pocket.

MD simulation rapidly reached dynamic equilibrium, with strong interactions between YFV_pro and c-myc remaining intact. However, some changes occurred at specific interaction sites (Fig. S12). SER 11, THR 12, PHE 40 formed multiple H-bonds with 5-terminal G-quartet plane and PHE 40 also formed *π*-*π* stacking with DG 13 which further facilitated the interaction between the residues and G-tetrad. Residues interacting with the loops changed into PHE 13, ARG 37, ALA 39 and LYS 47. More interestingly, the G4 flanking region (DT 1) penetrated more deeply into the binding pocket of the protein. It enhanced the stability of the G4-protein interaction, demonstrating the specificity of the recognition. The C-terminal region of the NS3 to which YFV_pro belongs functions as an RNA helicase, making it highly likely to participate in the recognition and binding of rG4 structures during viral RNA replication and translation. This interaction may facilitate its helicase activity, thereby promoting subsequent viral life processes.

The expressed YFV_pro is a serine protease (N-terminal part of NS3) composed of two *β*-barrels, each consisting of six antiparallel strands ^50^. In turn, these strands shape some pockets S1, S1’, S2, S3 and S4 (Fig. 4a), near the catalytic triad Ser-His-Asp, which is exceptionally important for accommodating side chains of substrates during recognition ^51^. In particular, the S2 pocket is created due to the existence of cofactor NS2B ^52^. It follows that NS2B is essential not only for the structure formation of the protease, but even for the realization of the function. To better understand the impact of G4 insertion into the protein binding pockets, we aligned and compared the crystal structure of the NS2B-NS3 protein complex (PDB: 6urv) with the structures obtained from our molecular docking and MD simulation (Fig. S13a & Fig. 4b). The results showed that the overall protein structure remained stable, with no changes observed in the interaction interface with G4. The only slight variation was in the region where NS2B was absent, leading to a minor increase in structural flexibility. From the surface mode (Fig. 4c), we could see that in the docking result, G4 flanking DT 1 only occupied S1’ pocket (Fig. S13b), while after MD simulation, the base shifted more deeply into the middle of S1 chamber (Fig. 4d).

**Figure 4.**
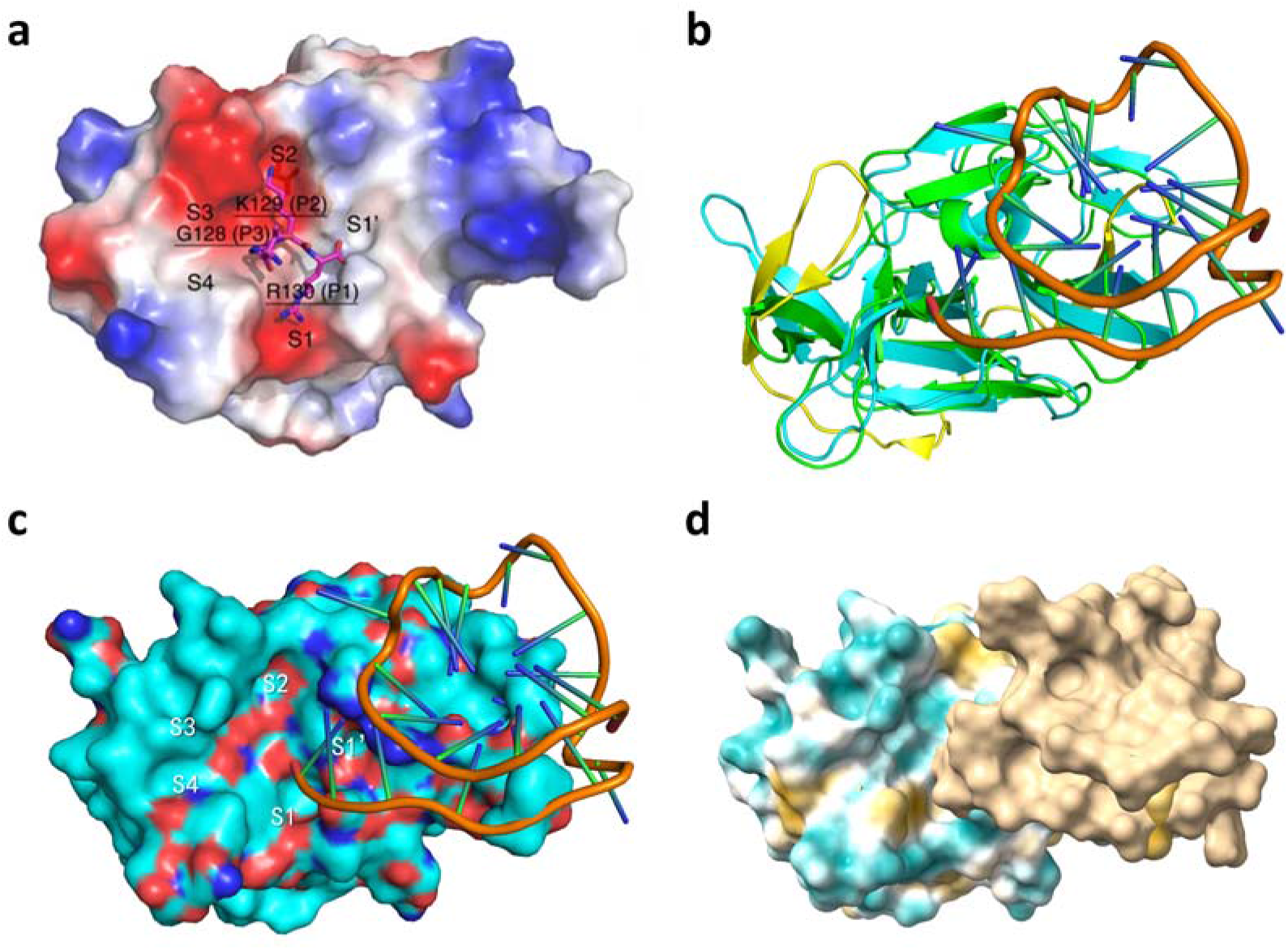
(**a**) Binding pockets of NS2B-NS3 complex with a small molecule. (**b**) Sticks mode of MD result aligned to NS2B-NS3 complex; cyan and yellow: NS3 and NS2B, green: NS3 of MD. (**c**) Surface mode of MD result aligned to NS2B-NS3 complex. (**d**) Surface mode of NS2B-NS3 complex with G4.

## Discussion

Among the candidate peptides identified by the FIMO method, our experiments revealed that some of them were indeed capable of G4 binding. Based on non-denaturing Electrophoresis and FRET-melting results, we considered the size and property of the original proteins, and eventually seleced two protein domain candidates (YEL and HAN), which are derived from Yellow Fever and Hantaan viruses.

We then discovered that the protein to which YEL belongs functions as both a protease and a helicase in Yellow Fever Virus, which suggests the potential for this protein to interact with rG4s during the viral cycle. HAN is derived from G1 and G2 glycoproteins in Hantaan. Since these glycoproteins form square-shaped surface spikes protruding from the viral membrane [32], they can be considered as promising targets for G4 aptamers. Thus, the interaction of both proteins with G4 structures offer interesting avenues for novel antiviral strategies.

During the expression of the two truncated proteins, we found that YFV_pro solubility in water was limited, consistent with reports that the flavivirus NS3 protein is not soluble alone *in vitro* and requires the assistance of NS2B for dissolution. Given that YFV_pro contains only the protease portion, we ended up aiding in its solubilization here by adding urea. Subsequent blotting experiments demonstrated that both DNA and RNA G4s bind to this protein. The FRET-melting results confirmed that G4 binds more strongly to YFV_pro than to its shorter 20-aa long peptide. The protein exhibits a preference for the parallel G4 topology. ThT competition assay and fluorescent anisotropy measurements confirmed binding and specificity. In contrast, HAN_pro performed poorly in our binding experiments. This may be explained by its three-dimensional structure (PDB 5LJX) (Fig. S3). For this reason, we focused our efforts on the YFV_pro.

The stable docking result of YFV_pro and c-myc indicates that the RGG matched motif indeed serves as the interaction site with G4. The α-helix stacks onto the G4 plane, forming H-bonds, electrostatic interactions and so on. Additionally, amino acid residues insert into the G4 lateral loops, and the flanking region is partially embedded within the protein structure. Subsequent MD showed some changes but maintained a relatively tight interaction state. Residues interacting with the G4 plane increased, and the appearance of π-π stacking further strengthened the interaction. The interaction between residues and G4 loops adjusted, indicating that a more stable interaction mode was established during the dynamic process. Such binding may also occur between rG4 formed during viral replication and NS3 protein, facilitating NS3-mediated unwinding of RNA secondary structures and ultimately promoting following viral processes. Additionally, the insertion at the flanking region became deeper, occupying the S1 protein pocket of the serine protease, which would significantly impact the catalytic activity of the enzyme ^53^. Exogenous G4 aptamer c-myc may inhibit this activity, possibly interfering with the cleavage of subsequent functional and structural proteins of the YFV virus, affecting its replication.

## Conclusion

Our study demonstrates that viruses causing hemorrhagic fevers harbor functional G4-binding proteins. By leveraging a conserved G4-binding motif, we identified candidate G4-binding peptides in several viruses and confirmed that the Yellow Fever virus NS3 protease binds to dG4/rG4 with high specificity and stability. Notably, this viral protein shows a distinct preference for parallel G4 conformations and engages specific RGG-motif residues, underscoring a previously unrecognized strategy by which viruses might interact with G4 structures during their life cycle. These findings mark a significant advance in our understanding of G4 – protein recognition, extending this concept to novel viral systems and revealing a new facet of viral molecular biology. By uncovering a viral enzyme that targets G4 elements, our work provides virology with a novel perspective on virus–host interplay and suggests that G4 structures could be exploited by viruses as part of their replication or regulatory mechanisms. This insight opens up new avenues for antiviral strategies – for example, designing G4-mimicking inhibitors or aptamers to selectively disrupt critical virus–G4 interactions.

It is important to note that our investigation focused on a select set of viral proteins and was conducted primarily *in vitro*; thus, the in vivo relevance of these G4 interactions remains to be established. Future studies should explore the prevalence and roles of G4-binding proteins across viruses and confirm their function during infection. Additionally, high-resolution structural studies (e.g., X-ray crystallography or cryo-EM) of viral protein–G4 complexes would provide deeper insight into the molecular basis of this interaction and guide the development of targeted therapeutics. Our work lays critical groundwork by expanding the scope of G4 biology to viruses causing hemorrhagic fevers. It underscores the potential of G4 structures as both fundamental elements in viral pathology and innovative targets for future antiviral interventions.

## Acknowledgments

This work has been conducted in the sustainability period of the project SYMBIT No. CZ.02.1.01/0.0/0.0/15_003/0000477 as its follow-up activity. The authors thank all their colleagues from the Laboratoire d’Optique & Biosciences for helpful discussions. J.L.M. and J.W. thank Laurent Lacroix (IBENS, Paris) for helpful discussions.

## Authors contributions

JW: Experimental design, biophysical and biochemical experiments, cloning, data analysis, writing the MS; VB: bioinformatics studies; RL: MD simulations; AC: biophysical characterization of G4 sequences, biochemical experiments; JLM: project design, funding acquisition, data analysis, writing the MS. All authors contributed to the editing of the MS.

