## Supplementary information for "Unraveling Viral peptide-G4 Interactions: the NS3 Protease Domain of Yellow Fever Virus Binds G-Quadruplexes with High Specificity and Affinity"

**Supplementary information – Jiawei Wang et al.**

**Supplementary Table S1:** Viruses causing viral hemorrhagic fevers (VHFs) considered here

| **Family** | **Genre** | **Name,**  **(Abbreviation)***^a^* **&**  **Virus acronym** | **Viral genome** |
| --- | --- | --- | --- |
| *Arenaviridae* | *Arenavirus* | Lassa (LAS)  LASV | ssRNA (*), bi-segmented, ≈7+3.4kb |
| *Bunyaviridae* | *Hantavirus* | Hantaan (HAN)  HTNV | ssRNA (-), tri-segmented, ≈6,5+3,7+2kb |
| *Filoviridae* | *Filovirus* | Ebola (EBO)  EBOV | ssRNA (-), non segmented, ≈19kb |
|  |  | Marburg (MAR)  MARV | ssRNA (-), non segmented, ≈19kb |
| *Flaviviridae* | *Flavivirus* | Yellow fever (YEL)  YFV | ssRNA (+), non segmented, 11-12 kb |
| *Paramyxoviridae* | *Henipahvirus* | Nipah (NIP)  NIV | ssRNA (-), non segmented, ≈18.2kb |
| *Nairoviridae* | *Orthonairovirus* | Crimea-Congo fever (CCH) CCHFV | ssRNA (-), tri-segmented, ≈11+4,4+1,7kb |

*^a^* : abbreviation used for the peptide originating from this virus. (*): ambisense.

**Supplementary Table S2.** YFV and HAN plasmids

| Acronym | Insertion site name | Vector name | Insert sequence |
| --- | --- | --- | --- |
| YFV | NdeI_XhoI | pET-28a(+) | TTGGAGGATGGCATTTATGGCATCTTTCAAAGCACATTTTTGGGTGCGTCGCAGCGTGGTGTTGGCGTGGCGCAAGGAGGTGTATTTCATACAATGTGGCATGTTACACGTGGAGCTTTTCTGGTTCGTAATGGTAAAAAACTGATACCATCTTGGGCGAGCGTTAAAGAAGATCTGGTTGCGTATGGTGGTAGTTGGAAGCTGGAAGGTCGCTGGGATGGCGAAGAAGAAGTCCAATTAATAGCCGCCGTGCCCGGTAAGAACGTAGTAAATGTACAAACTAAGCCGAGTCTGTTCAAGGTACGTAACGGAGGGGAAATTGGGGCAGTTGCGCTGGATTATCCAAGCGGCACCAGTGGTTCTCCGATCGTTAATCGAAATGGCGAAGTGATTGGGTTATATGGAAACGGCATCCTGGTTGGGGACAACTCGTTTGTTTCAGCAATCTCTCAGACATGA |
| HAN | NdeI_XhoI | pET-28a(+) | AACCTGGGTGAGAACCCGTGCAAAATCGGCTTACAAACCTCATCCATTGAGGGCGCCTGGGGTAGCGGTGTGGGGTTTACGCTGACATGCCTCATTTCGCTGACTGAATGCCCTACATTTCTCACATCAATAAAAGCATGTGACAAAGCCATTTGTTATGGCGCCGAAAGTGTTACGCTTACTCGCGGGCAGAATACTGTGAAGGTTTCAGGTAAGGGTGGCCACTCCGGTAGTTCCTTTAAATGTTGCCATGGGGAGGATTGCTCTCAAATTGGTCTCCATGCGGCAGCTCCACATCTTTAG |

**Supplementary Table S3.** Matched sequence of NIQI motif retrieved by FIMO approach

| **Virus** | **Alt ID** | **Sequence Name** | **p-value** | **Matched Sequence** |
| --- | --- | --- | --- | --- |
| **CCHFV** | **RGRGRGRGGGSGGSGGRGRG** | **lcl\|NC_005301.3_prot_YP_325663.1_1** | **3.16E-04** | **RYRLCSKGGVEQHSEEDLRR** |
| **Ebola** | **RGRGRGRGGGSGGSGGRGRG** | **lcl\|NC_002549.1_prot_NP_066247.1_5** | **2.66E-04** | **RGFPRCRYVHKVSGTGPCAG** |
| **Ebola** | RGRGRGRGGGSGGSGGRGRG | lcl\|NC_002549.1_prot_NP_066248.1_6 | 2.66E-04 | RGFPRCRYVHKVSGTGPCAG |
| **Ebola** | RGRGRGRGGGSGGSGGRGRG | lcl\|NC_002549.1_prot_NP_066246.1_4 | 2.66E-04 | RGFPRCRYVHKVSGTGPCAG |
| **Hantaan** | **RGRGRGRGGGSGGSGGRGRG** | **AFM85164.1** | **1.68E-04** | **TVKVSGKGGHSGSSFKCCHG** |
| **Hantaan** | RGRGRGRGGGSGGSGGRGRG | AFM85165.1 | 1.82E-04 | MLTTRGRQTTKDNKGTRIRF |
| **Hantaan** | RGRGRGRGGGSGGSGGRGRG | AFM85164.1 | 8.57E-04 | RGQNTVKVSGKGGHSGSSFK |
| **Lassa** | **RGRGRGRGGGSGGSGGRGRG** | **lcl\|NC_004296.1_prot_NP_694870.1_2** | **1.02E-04** | **QTFMRMAWGGSYIALDSGRG** |
| **Lassa** | RGRGRGRGGGSGGSGGRGRG | lcl\|NC_004296.1_prot_NP_694869.1_1 | 1.48E-04 | RRALLNMIGMSGGNQGARAG |
| **Lassa** | RGRGRGRGGGSGGSGGRGRG | lcl\|NC_004296.1_prot_NP_694869.1_1 | 8.57E-04 | IGMSGGNQGARAGRDGVVRV |
| **Marburg** | **RGRGRGRGGGSGGSGGRGRG** | **lcl\|JX458836.1_prot_AFV31217.1_5** | **1.87E-04** | **RGRSRTRNHQAIPSIYHETQ** |
| **Marburg** | RGRGRGRGGGSGGSGGRGRG | lcl\|JX458836.1_prot_AFV31219.1_7 | 7.00E-04 | LARRIKGQRGSLRSNWRFIG |
| **Nipah** | **RGRGRGRGGGSGGSGGRGRG** | **lcl\|NC_002728.1_prot_NP_112027.1_8** | **3.74E-05** | **RGVSKQRIIGVGEVLDRGDE** |
| **Nipah** | RGRGRGRGGGSGGSGGRGRG | lcl\|NC_002728.1_prot_NP_112025.1_6 | 7.00E-04 | RKIDRMKLQFSLGSIGGLSL |
| **Yellow fever** | **RGRGRGRGGGSGGSGGRGRG** | **lcl\|NC_002031.1_prot_NP_041726.1_1** | **7.34E-05** | **RGVGVAQGGVFHTMWHVTRG** |
| **Yellow fever** | RGRGRGRGGGSGGSGGRGRG | lcl\|NC_002031.1_prot_NP_041726.1_1 | **7.34E-05** | HERGYVKLEGRVIDLGCGRG |
| **Yellow fever** | RGRGRGRGGGSGGSGGRGRG | lcl\|NC_002031.1_prot_NP_041726.1_1 | 1.23E-04 | SDRGWGNGCGLFGKGSIVAC |
| **Yellow fever** | RGRGRGRGGGSGGSGGRGRG | lcl\|NC_002031.1_prot_NP_041726.1_1 | 1.48E-04 | RGWGNGCGLFGKGSIVACAK |
| **Yellow fever** | RGRGRGRGGGSGGSGGRGRG | lcl\|NC_002031.1_prot_NP_041726.1_1 | 3.16E-04 | AIKGPLRISASSAAQRRGRI |
| **Yellow fever** | RGRGRGRGGGSGGSGGRGRG | lcl\|NC_002031.1_prot_NP_041726.1_1 | 5.01E-04 | VGRGDSRLTYQWHKEGSSIG |
| **Yellow fever** | RGRGRGRGGGSGGSGGRGRG | lcl\|NC_002031.1_prot_NP_041726.1_1 | 5.01E-04 | RGYVKLEGRVIDLGCGRGGW |
| **Yellow fever** | RGRGRGRGGGSGGSGGRGRG | lcl\|NC_002031.1_prot_NP_041726.1_1 | 6.73E-04 | LVFSPGRKNGSFIIDGKSRK |
| **Yellow fever** | RGRGRGRGGGSGGSGGRGRG | lcl\|NC_002031.1_prot_NP_041726.1_1 | 8.57E-04 | CKRTYSDRGWGNGCGLFGKG |
| **Yellow fever** | RGRGRGRGGGSGGSGGRGRG | lcl\|NC_002031.1_prot_NP_041726.1_1 | 8.57E-04 | TYSDRGWGNGCGLFGKGSIV |
| **Yellow fever** | RGRGRGRGGGSGGSGGRGRG | lcl\|NC_002031.1_prot_NP_041726.1_1 | 8.57E-04 | EGRVIDLGCGRGGWCYYAAA |

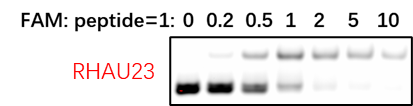

**Supplementary Figure S1.** RHAU23 (HPGHLKGREIGMWYAKKQGQKNK) is a G4-binding peptide; a retarded band is observed upon peptide addition.

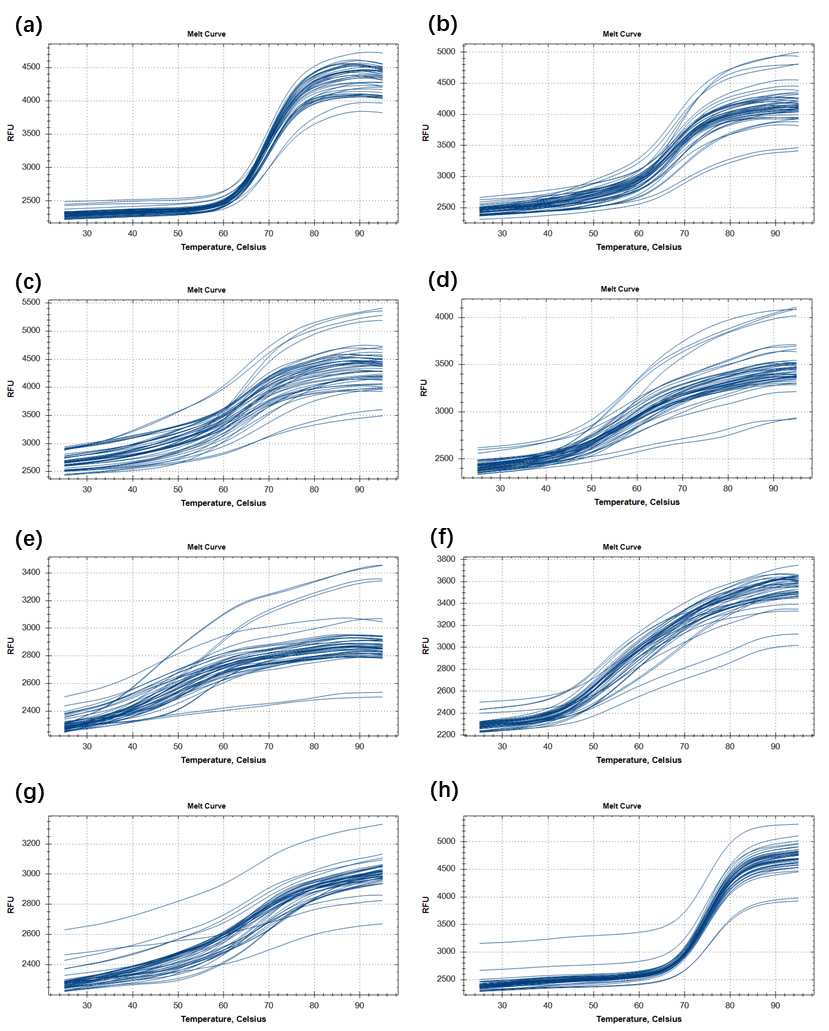

**Supplementary Figure S2.** Melting curves of peptides with (a) FdxT, (b) F25CebT, (c) FmycT, (d) F21CTAT, (e) FTBAT, (f) F21T, (g) FR21T, (h) FRdxT.

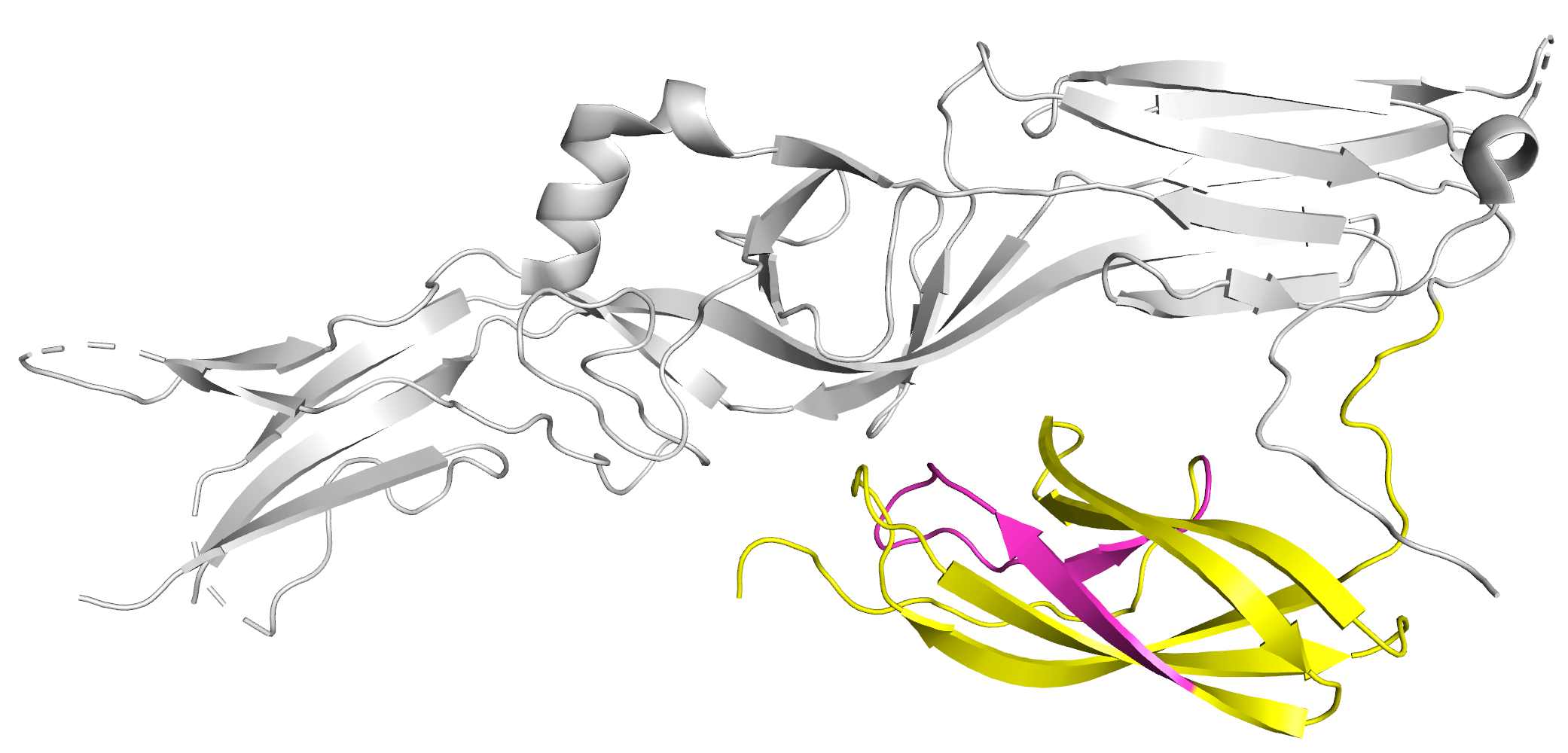

**Supplementary Figure S3.** Intercepted 100-aa peptide HAN_pro (yellow) in 5ljx; RGG matched motif (pink) in HAN_pro.

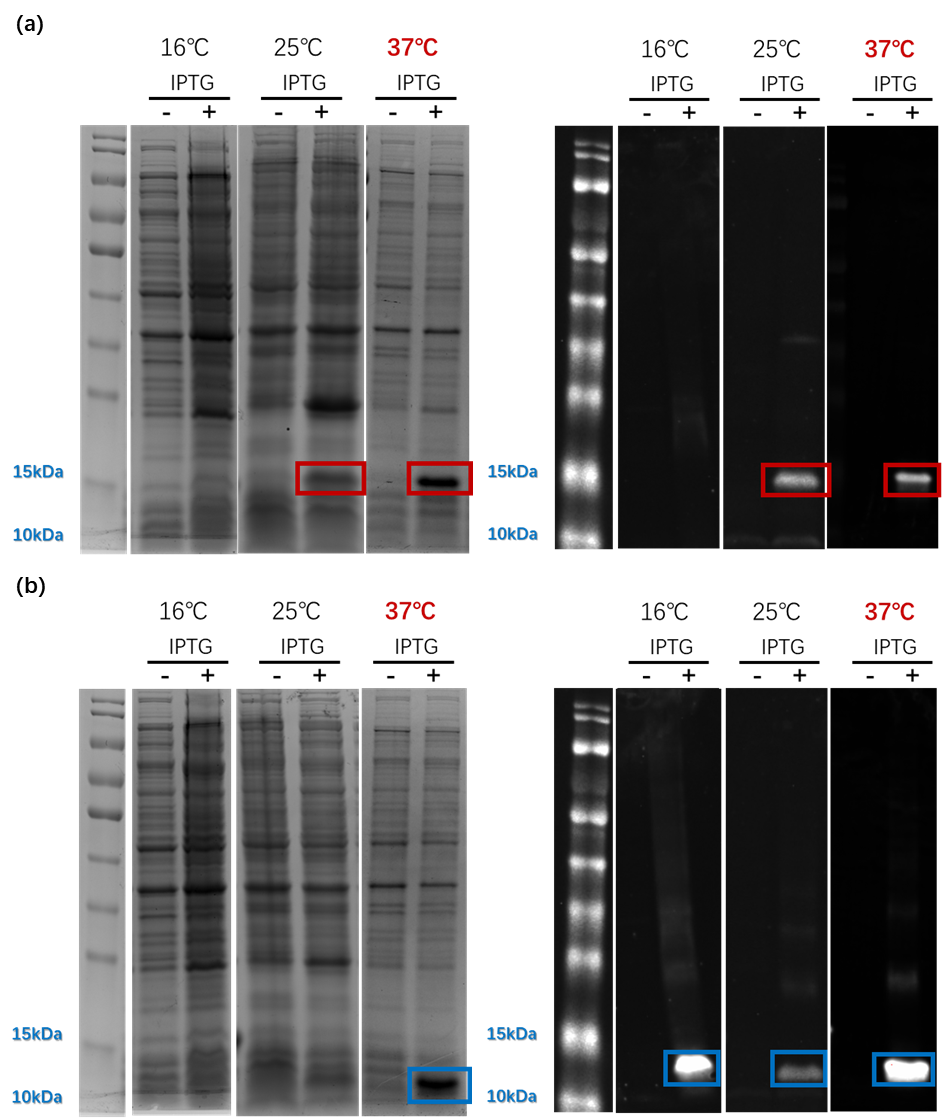

**Supplementary Figure S4.** Cultures under different temperatures for (a) YFV_pro and (b) HAN_pro.

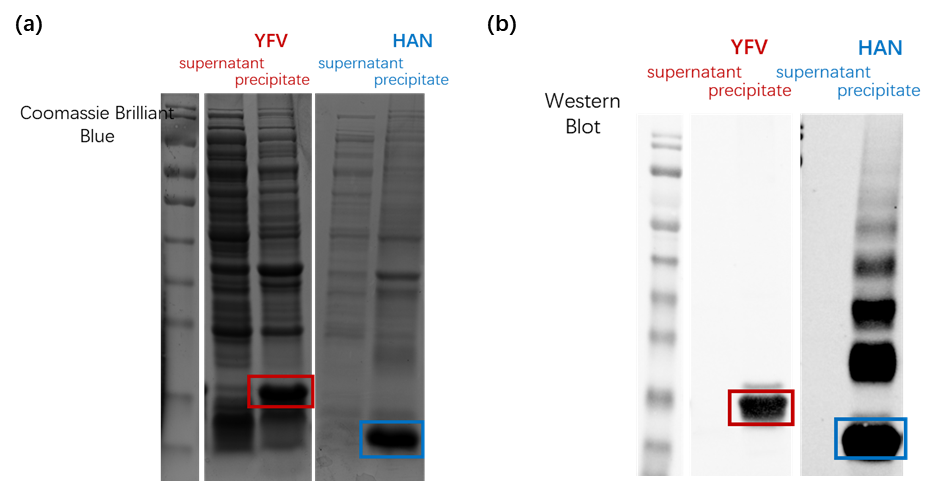

**Supplementary Figure S5.** Protein expression quantity of different components

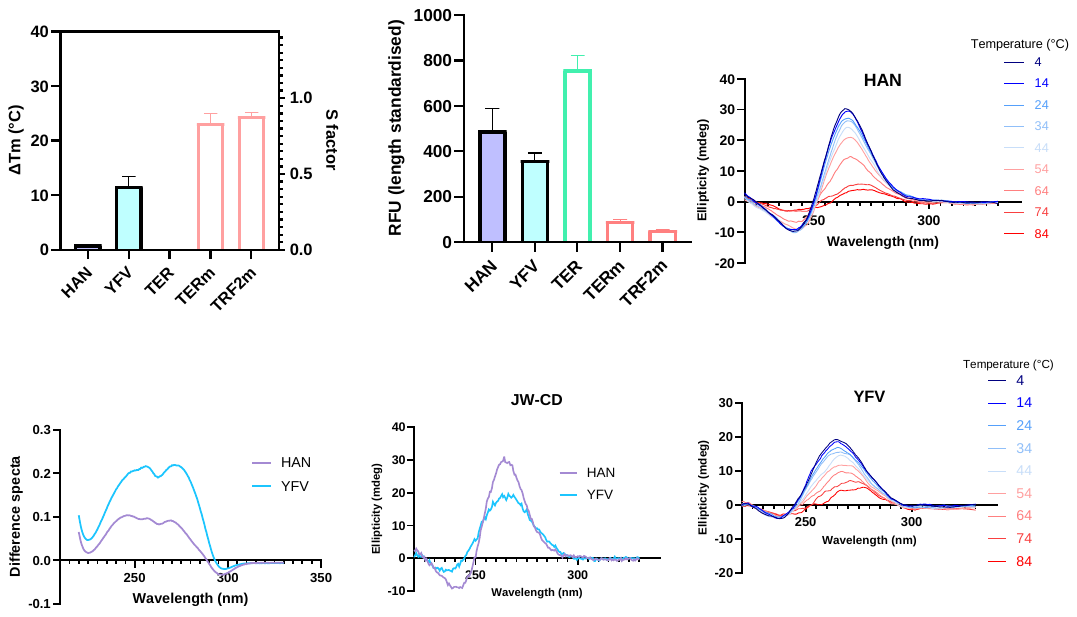

**Supplementary Figure S6.** Annealing of rG4 YFV_G4 and HAN_G4 on CD.

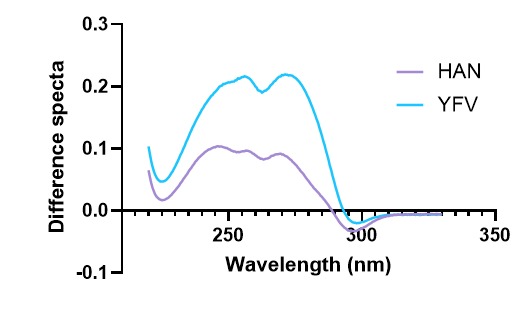

**Supplementary Figure S7.** TDS for rG4 YFV_G4 and HAN_G4.

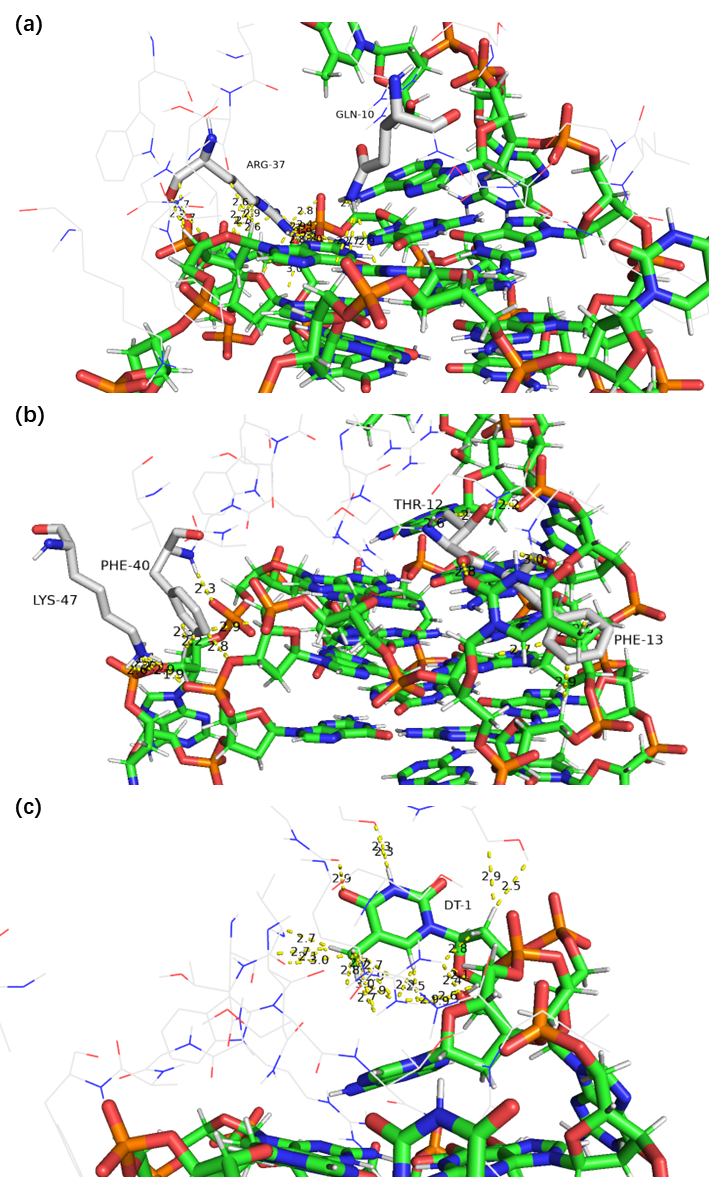

**Supplementary Figure S8.** Docking model. (a) Interactions on G-tetrad; (b) Interactions with loops; (c) Insertion of flanking. Green sticks: G4; White sticks: amino acid residues.

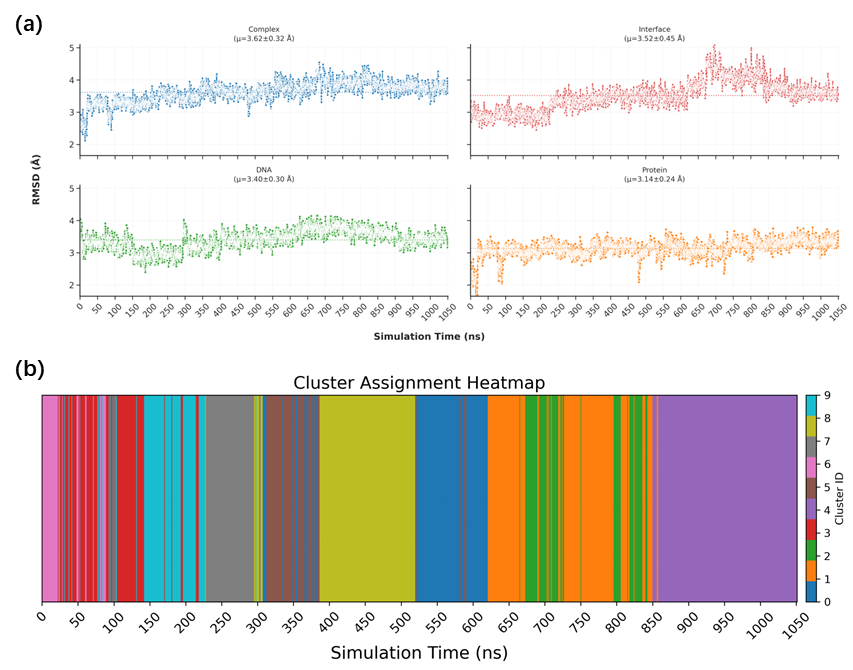

**Supplementary Figure S9.** Judgement parameters of MD result.

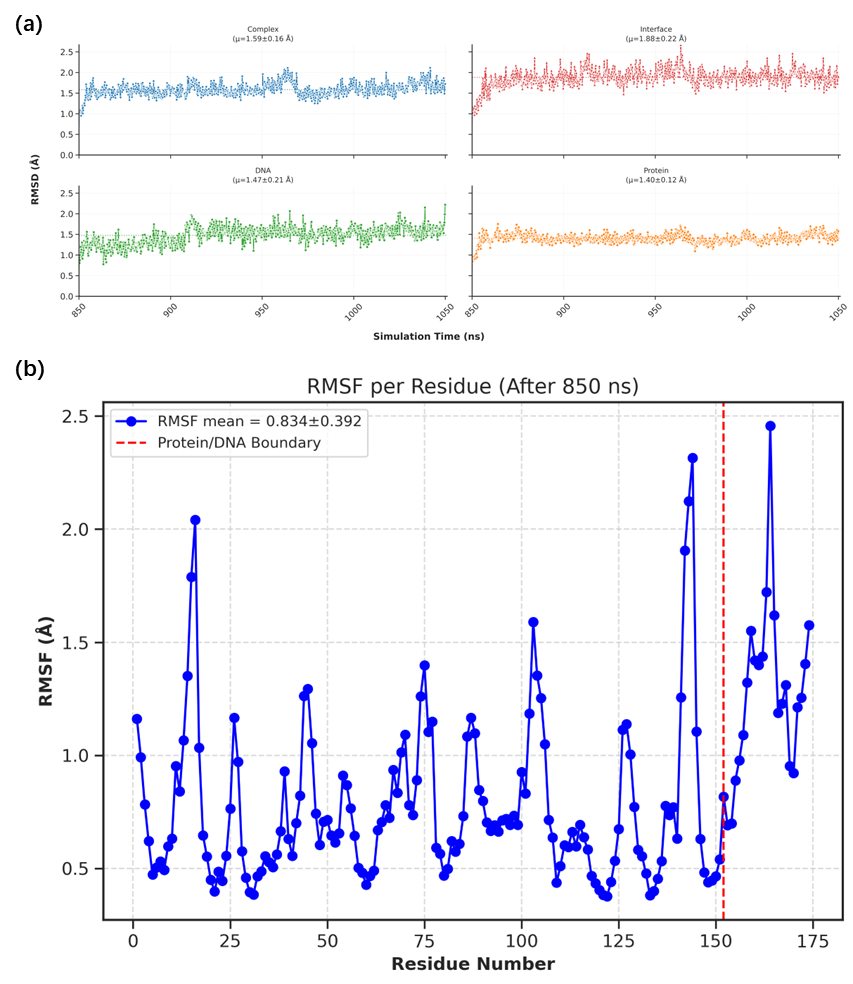

**Supplementary Figure S10.** RMSD and RMSF after 850ns.

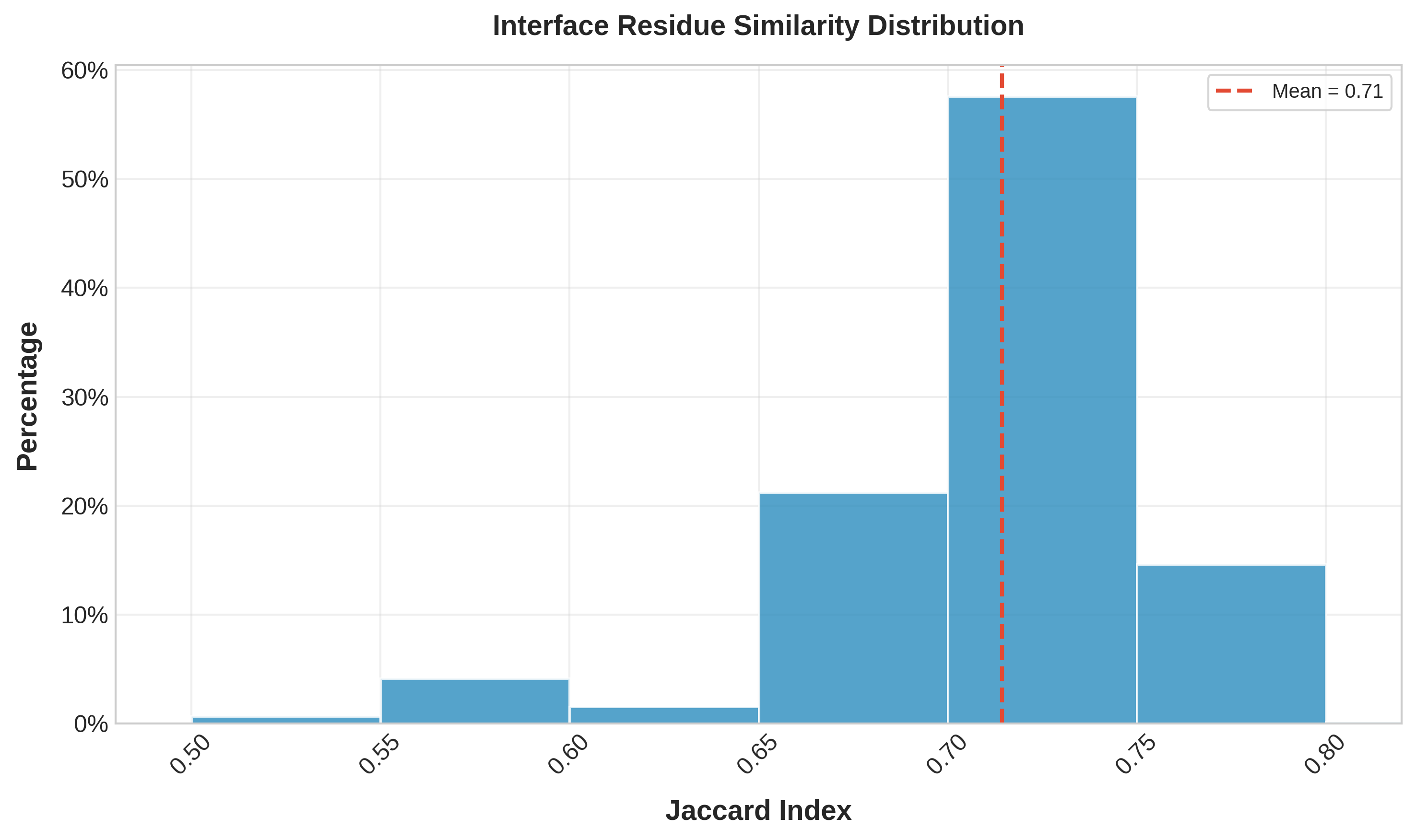

**Supplementary Figure S11.** The Jaccard index of the interface.

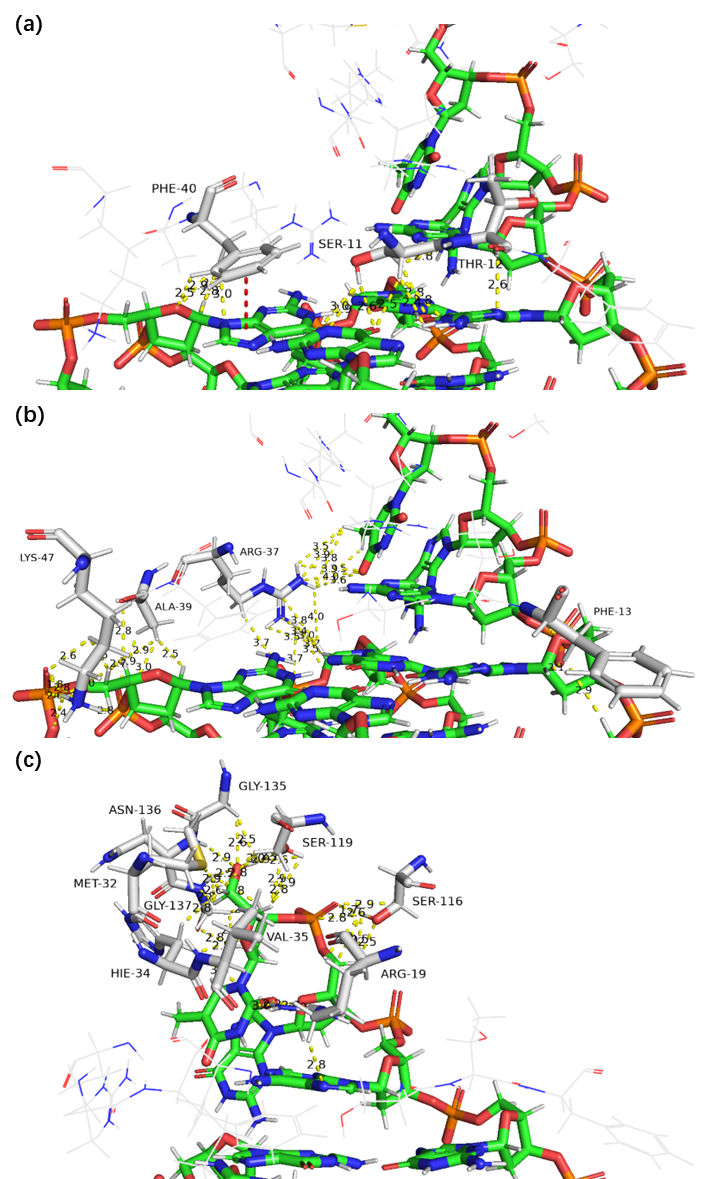

**Supplementary Figure S12.** MD simulation model. (a) Interactions on G-tetrad; (b) Interactions with loops; (c) Insertion of flanking. Green sticks: G4; White sticks: amino acid residues.

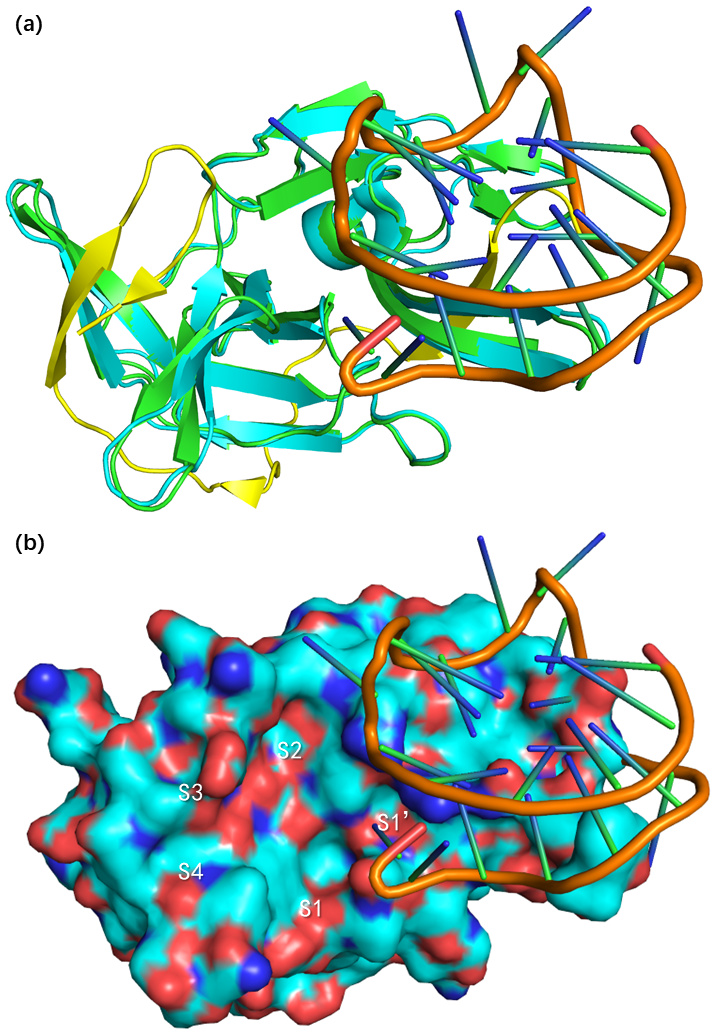

**Supplementary Figure S13.** (a) Sticks mode of docking result aligned to NS2B-NS3 complex; cyan and yellow: NS3 and NS2B, green: NS3 of docking. (b)Surface mode of docking result aligned to NS2B-NS3 complex.
